# Super-resolved, three-dimensional spatial transcriptomics reveals cell-type and brain-region-specific modulation of key epitranscriptomic switches following adolescent alcohol exposure

**DOI:** 10.64898/2026.01.22.698893

**Authors:** Jamuna Tandukar, Anam Islam, Emir Malovic, Bisma Afzal, Huaibo Zhang, Subhash C. Pandey, Ruixuan Gao

## Abstract

Epitranscriptomic mechanisms dynamically regulate neuronal function through gene expression, but their precise roles in neuropsychiatric and neurological disorders remain to be fully elucidated. A major obstacle to advancing such studies is the absence of a methodology for precise, cell-type and brain-region-specific quantification of critical epitranscriptomic regulators under these complex brain conditions. To overcome this challenge, we developed a super-resolved, three-dimensional spatial transcriptomics method to quantify key epitranscriptomic switches in intact brains. Using this method, we quantified the expression of *Mettl3*, an N6-methyladenosine (m6A) methyltransferase enzyme recently shown to be upregulated in the amygdala after adolescent intermittent ethanol (AIE) exposure in rats. We observed a significant increase in cytoplasmic *Mettl3* mRNA in neurons, but not in astrocytes or microglia, within the adult central amygdala and the CA1 and dentate gyrus of hippocampus following AIE. In contrast, no significant changes were observed across neurons, astrocytes, or microglia within the basolateral amygdala or the hippocampal CA3.

Additionally, we found both the cytoplasmic density and subcellular localization of *Mettl3* mRNA were dependent on the specific cell types and brain subregions examined. These results suggest that AIE increases *Mettl3* expression in a highly cell-type-specific and spatially heterogeneous manner, underscoring the necessity of high-resolution spatial transcriptomics methods for studying transcriptomic and epitranscriptomic regulations under neurological conditions.

**Significance Statement:** Epitranscriptomics plays a crucial role in neuronal functions by influencing the splicing, stability, and translation of genes. However, the exact role of epitranscriptomic mechanisms, such as m6A RNA methylation, in brain disorders remains unclear, particularly in a cell-type and circuitry-specific manner. Here we developed a super-resolved, three-dimensional spatial transcriptomics method and applied it to a model of alcohol exposure. We found differential cell-type- and brain-region-specific modulation of *Mettl3*, a key m6A enzymatic switch, across major brain regions following adolescent intermittent ethanol exposure in adulthood. Our findings, coupled with our pipeline, are expected to address existing methodological limitations and knowledge gaps, thereby accelerating brain transcriptomic and epitranscriptomic studies under various psychiatric and neurological conditions.

## Introduction

Epitranscriptomics plays a crucial and increasingly recognized role in neurobiology, influencing a broad range of processes from neural development and plasticity to learning, memory, and the pathogenesis of neurological disorders (Shafik et al., 2022; Malovic and Pandey, 2023; Zhang et al., 2024). The brain exhibits a particularly rich and dynamic epitranscriptome, with modifications like N6-methyladenosine (m6A) being highly abundant in neuronal cells (He and He, 2021; Yu et al., 2021; Zhang et al., 2024). Recently, m6A modification has been shown to regulate neurogenesis, axonal guidance, and regeneration, as well as synaptic function by affecting mRNA splicing, stability, and translation of crucial neuronal genes (Yu et al., 2021; Castro-Hernández et al., 2023).

Alcohol use and misuse are major dietary and environmental factors that trigger functional, behavioral, and anatomical changes in the brain (Koob and Volkow, 2016; Spindler et al., 2021). Previous studies have shown that early-life or adult ethanol exposure causes lasting circuit, cellular, and molecular modifications in brain regions controlling reward, stress, emotions, cognition, and decision-making, including the prefrontal cortex, amygdala, hippocampus, and striatum (Oscar-Berman and Marinković, 2007; Koob and Volkow, 2016; Pandey et al., 2017). Notably, recent studies have suggested that epigenetic mechanisms play a crucial role in mediating these changes, thereby establishing alcohol exposure and addiction as an ideal model for epigenetic and epitranscriptomic studies in the brain (Pandey et al., 2015, 2017; Bohnsack et al., 2022; Maccioni et al., 2025; Malovic et al., 2025). Indeed, our recent study has found widespread and significant changes in key m6A modifiers in the amygdala, medial prefrontal cortex, hippocampus, and nucleus accumbens during adolescence or adulthood following adolescent intermittent ethanol (AIE) exposure. Specifically, methyltransferase-like 3 (METTL3), the enzymatic subunit of the m6A methyltransferase complex, was upregulated in the amygdala during adulthood after AIE exposure, and pharmacological inhibition of METTL3 attenuated the AIE-induced anxiety-like behaviors (Malovic et al., 2025). However, few studies to date have explored alcohol-induced epitranscriptomic changes in intact brains in a cell-type-specific manner, particularly in the amygdala or other relevant brain regions.

One factor contributing to the lack of such investigations is the absence of a reliable and accessible approach that links system-level physiological and behavioral perturbations with cellular and molecular epitranscriptomic changes in a cell-type and brain-region-specific manner. From a methodological perspective, single-cell transcriptomics and epitranscriptomics have made tremendous advances in understanding tissue-wide heterogeneity of gene expression and associated epitranscriptomic changes (Crespo-García et al., 2024), but lack spatial information or resolution to study specific cells within a particular brain subregion. Recent development and commercialization of spatial transcriptomics methods have advanced the capability of capturing gene expression patterns in intact brain tissue. However, these technologies are often limited in their ability to achieve both cellular resolution and accurate quantification. For instance, sequencing-based approaches (Ståhl et al., 2016; Rodriques et al., 2019; Bressan et al., 2023) are powerful tools for quantifying RNA transcripts across large tissue areas but lack subcellular resolution. Imaging-based techniques (Lubeck et al., 2014; Chen et al., 2015; Eng et al., 2019) can achieve single-molecule detection with subcellular resolution, but are often constrained by signal crowding and optical diffraction, particularly in densely-populated brain tissue, complicating accurate assignments of transcripts to individual cells and subcellular compartments (Chen et al., 2015; Eng et al., 2019; Xia et al., 2019).

The recent development of expansion microscopy has addressed a number of these limitations by spatially decrowding RNA molecules in intact brain tissue and enabling molecular mapping at effective super-resolution (Wang et al., 2018, 2021; Alon et al., 2021; Cui et al., 2023; Sarfatis et al., 2025). Given these advances, we hypothesized that by combining brain tissue expansion with highly sensitive RNA labeling and detection, we can accurately quantify key epitranscriptomic switches at cell-type and brain-region specific manner after alcohol exposure. To demonstrate this, we subjected rats to AIE exposure and applied a tissue-expansion-based spatial transcriptomics approach to investigate changes in the mRNA expression of METTL3 in the brain. Using our pipeline, we confirmed that AIE exposure significantly increases *Mettl3* mRNA levels within neurons of discrete amygdalar and hippocampal subregions, while adjacent subregions and non-neuronal cell types remain unaffected. We expect our findings, along with the super-resolved, three-dimensional spatial transcriptomic pipeline developed here, will bridge the methodological and knowledge gaps in brain epitranscriptomic and transcriptomic studies.

## Materials and Methods

### Animals

All animal experiments were conducted in accordance with the National Institutes of Health Guide for the Care and Use of Laboratory Animals and approved by the IACUC at the University of Illinois Chicago. Sprague-Dawley rats were purchased from Envigo RMS (Indianapolis, Indiana) at postnatal day 17 with their respective dams. The rats were housed in the standard 12-hour light and dark cycle with ad libitum access to food and water, as previously described (Malovic et al., 2025).

### Experimental design

Alcohol was administered following previously described methods (Malovic et al., 2025). In brief, 12 animals were used for this study, 3 female and 3 male rats per experimental group. Rat pups were weaned on postnatal day 21 and pair housed. Adolescent intermittent saline (AIS) or ethanol (AIE) intraperitoneal administration began on postnatal day 28 and completed on postnatal day 41. Rats were given 2 g/kg (20% w/v) of ethanol or saline in a 2-day “on” and 2-day “off” schedule (totaling 8 injection days from postnatal day 28 to 41). The rats were sacrificed at postnatal day 95 under isoflurane anesthesia, followed by 4% paraformaldehyde perfusion in nuclease-free conditions. The whole brains were collected and stored in nuclease-free 1x PBS (AM9625, ThermoFisher) at 4°C until sectioning.

### Tissue preparation

The brains were sliced into ∼100 µm coronal brain sections using a vibratome (Leica VT1200S). The collected brain sections (3 brain sections per animal) were stored in RNA*later* solution (AM7020, ThermoFisher) at 4°C until further processing.

### RNA labeling

To label RNA, we utilized Hybridization Chain Reaction RNA fluorescence *in situ* hybridization v3.0 (HCR RNA-FISH v3) (Choi et al., 2018). Briefly, brain sections were first permeabilized with 0.5% Triton X-100 in nuclease-free 1x PBS for 1 hour at room temperature (RT) and then washed for 30 minutes at RT using a wash buffer composed of 20% (v/v) formamide (17899, ThermoFisher) and 2x SSC (AM9770, ThermoFisher). Meanwhile, the hybridization buffer [10% (w/v) dextran sulfate (S4031, MilliporeSigma), 2x SSC, 10% (v/v) formamide] was preheated to 37°C. The brain sections were then incubated in the warmed hybridization buffer for 30 minutes at 37°C to facilitate pre-hybridization.

For probe hybridization, HCR RNA-FISH probes for *Mettl3* mRNA (Accession #: NM_001024794.1, Molecular Instruments, Inc.) and 28S ribosomal RNA (rRNA) (Accession #: NR_046246.3, Molecular Instruments, Inc.) were diluted in the preheated hybridization buffer to a final concentration of 32 nM and 48 nM, respectively. The brain sections were incubated in the probe hybridization buffer overnight on a shaker at 37°C. Next day, excess probes were removed by washing the brain sections with the wash buffer twice, 30 minutes each at 37°C. This was followed by sequential washes in 1x PBS, first for 2 hours at 37°C and then for another 2 hours at RT.

To prepare for amplification, the brain sections were incubated in an amplification buffer [10% (w/v) dextran sulfate, 5x SSC, 0.1% (v/v) Tween-20 (1610781, Bio-Rad)] for 30 minutes at RT. In parallel, fluorescently labeled HCR hairpins (B1 labeled with Alexa Fluor 488, B3 labeled with Alex Fluor 594, Molecular Instruments, Inc.) were snap-cooled by heating to 95°C for 90 seconds, then allowed to cool at RT in the dark for 30 minutes. Once prepared, the hairpins were mixed with the amplification buffer and immediately applied to the brain sections. Amplification was allowed to occur overnight on a shaker at 4°C. The following day, amplification was stopped with four washes with 5x SSCT [5x SSC, 0.1% (v/v) Tween-20], 30 minutes each at RT.

### Immunostaining

After the HCR RNA-FISH steps, the brain sections were incubated in a blocking buffer containing 1% nuclease-free BSA (AM2616, ThermoFisher) in 1x PBS with 0.1% Triton X-100 for 2 hours. Next, the brain sections were incubated in the primary antibody solution (1:200 dilution with the blocking buffer) overnight at RT. Primary antibodies against NeuN [guinea pig, 266004, Synaptic Systems (SYSY)], GFAP (guinea pig, 173308, SYSY), or IBA1 (chicken, 234009, SYSY) were used to label cells of each cell type in the sample (**Supplemental Table S1**). After washing out unbound primary antibodies with the blocking buffer three times, 10 minutes per wash, brain sections were incubated in the secondary antibody solution (1:200 dilution with the blocking buffer) overnight at RT. Secondary antibodies were prepared in house by conjugating 647 fluorescent dyes (SeTau-647-NHS-1mg, K9-4149, SETA BioMedicals) with unconjugated secondary antibodies (anti-guinea pig, A18771 or anti-chicken, A16056, ThermoFisher) (**Supplemental Table S2**). The following day, the brain sections were washed three times, 30 minutes each with the blocking buffer.

### Anchoring, gelation, and expansion

The RNA-labeled and immunostained brain sections were pre-incubated twice with 100 mM sodium bicarbonate (pH 8.5) for 15 minutes at RT. The brain sections were then treated with the anchoring reagent glycidyl methacrylate (GMA) (779342, Millipore Sigma) dissolved in 100 mM sodium bicarbonate for 3 hours at 37°C (Cui et al., 2023). To eliminate unreacted GMA, the brain sections were washed three times with nuclease-free 1x PBS, 5 minutes per wash.

Next, the brain sections were incubated in a gelling solution [1x PBS, 2 M NaCl, 8.625% (w/v) sodium acrylate, 2.5% (w/v) acrylamide, 0.15% (w/v) N,N′-methylenebisacrylamide, 0.01% (w/v) 4-hydroxy-2,2,6,6-tetramethylpiperidin-1-oxyl (4HT), 0.2% (w/v) ammonium persulfate (APS), and 0.2% (w/v) tetramethylethylenediamine (TEMED)] at 4°C for 20 minutes and then transferred to a humidified incubator at 37°C, where they were allowed to polymerize for 1.5 hours. The gelled brain sections were immediately immersed in proteinase K (proK) digestion buffer (1:100 dilution) (P8107S, New England Biolabs) and incubated overnight at RT for sample homogenization. Before expansion, the homogenized brain sections were stained with 10 µg/mL DAPI in nuclease-free 1x PBS for 30 minutes. Finally, the brain sections underwent three washes with nuclease-free water containing 0.05x SSC, 10 minutes per wash, to complete expansion.

### Confocal microscopy

The expanded brain sections were transferred to poly-L-lysine (A-005-C, MilliporeSigma)-coated imaging well plates. All images of the expanded brain sections were captured using a Nikon spinning disk confocal microscope (CSU-W1, Yokogawa) with a 40x 1.15 NA water-immersion objective (Nikon), controlled with NIS-Elements AR imaging software v5.30.04 (Nikon). Four color channels (405 nm, 488 nm, 561 nm, and 640 nm) were used for imaging.

### Three-dimensional (3D) RNA puncta quantification

To ensure unbiased analysis, images were processed in a randomized and blind manner. Briefly, the AIS and AIE images were re-labeled and randomized. The 488 nm channel containing *Mettl3* mRNA signals was excluded prior to cell selection and segmentation. A researcher blind to the experimental conditions and original file labels randomly cropped two to three cells from each field of view per cell type using the 405 nm (DAPI), 561 nm (28S rRNA), and 640 nm (NEUN, GFAP, or IBA1) channels.

The resulting image stacks encompassing individual cells were drift-corrected using the Correct 3D Drift plugin in ImageJ (version 1.54f), when necessary. 3D segmentation of the cytoplasm and the subsequent quantification of cytoplasmic mRNA puncta were performed using a customized pipeline in CellProfiler (version 4.2.6) that utilized its built-in modules for 3D analysis. Briefly, adaptive thresholding was applied to identify cytoplasmic contours above the image background. Watershed module was used to segment cells into individual objects, and size-based filtering was applied to remove non-target masks. For *Mettl3* puncta, adaptive thresholding and size-filtering was applied to identify puncta above the image background and to remove noise and false positives. Finally, *Mettl3* puncta were counted within each segmented cell mask using the RelateObjects module. For quantification of *Mettl3* RNA in the nucleus, nuclear masks were generated from the DAPI label, similar to the generation of cytoplasmic masks.

For neurons and astrocytes, the 28S rRNA label was used for cytoplasmic segmentation. For microglia, the IBA1 label was used for cytoplasmic segmentation because the rRNA signals were not a reliable cytoplasmic label. Briefly, cell-body masks were generated from the IBA1 label, using the same method as the cytoplasmic mask generation. The final cytoplasmic puncta count was determined by subtracting the puncta counted within the nuclear mask from the total count within the cell-body mask.

Cytoplasmic volumes were measured using the abovementioned segmentation pipeline and were subsequently converted to the pre-expansion biological scale.

Cytoplasmic *Mettl3* puncta density was calculated by dividing the number of cytoplasmic *Mettl3* puncta by the corresponding cytoplasmic volume.

## Visualization

All 3D renderings were generated using Imaris (version 10.1.1, Oxford Instruments). A gamma (γ) adjustment of 1.3 was applied to the 561 nm channel (rRNA).

## Statistical analysis

All statistical analysis was performed using OriginLab (version 2024b). The statistical tests and the sample numbers used for each experiment are indicated in the corresponding figure captions. Unless mentioned, all groups were subjected to two-tailed t-test with post-hoc Welch correction.

## Results

### RESOLution enhanced Visualization using Expansion-coupled FISH (RESOLVE-FISH) enables super-resolved, three-dimensional spatial transcriptomics in intact brain tissue

To enable accurate quantification of target biomolecules, including cytoplasmic *Mettl3* mRNA, at the single-cell level, we developed RESOLution enhanced Visualization using

Expansion-coupled FISH (RESOLVE-FISH), an experimental and computational pipeline that combines tissue expansion, highly sensitive molecular labeling, and post-imaging single-cell analysis (**Fig. 1**). By integrating cell-type markers that provide precise single-cell identification in the post-imaging analysis, RESOLVE-FISH allows accurate assignments of fluorescently labeled target molecules to specific cellular and subcellular compartments.

**Fig. 1:**
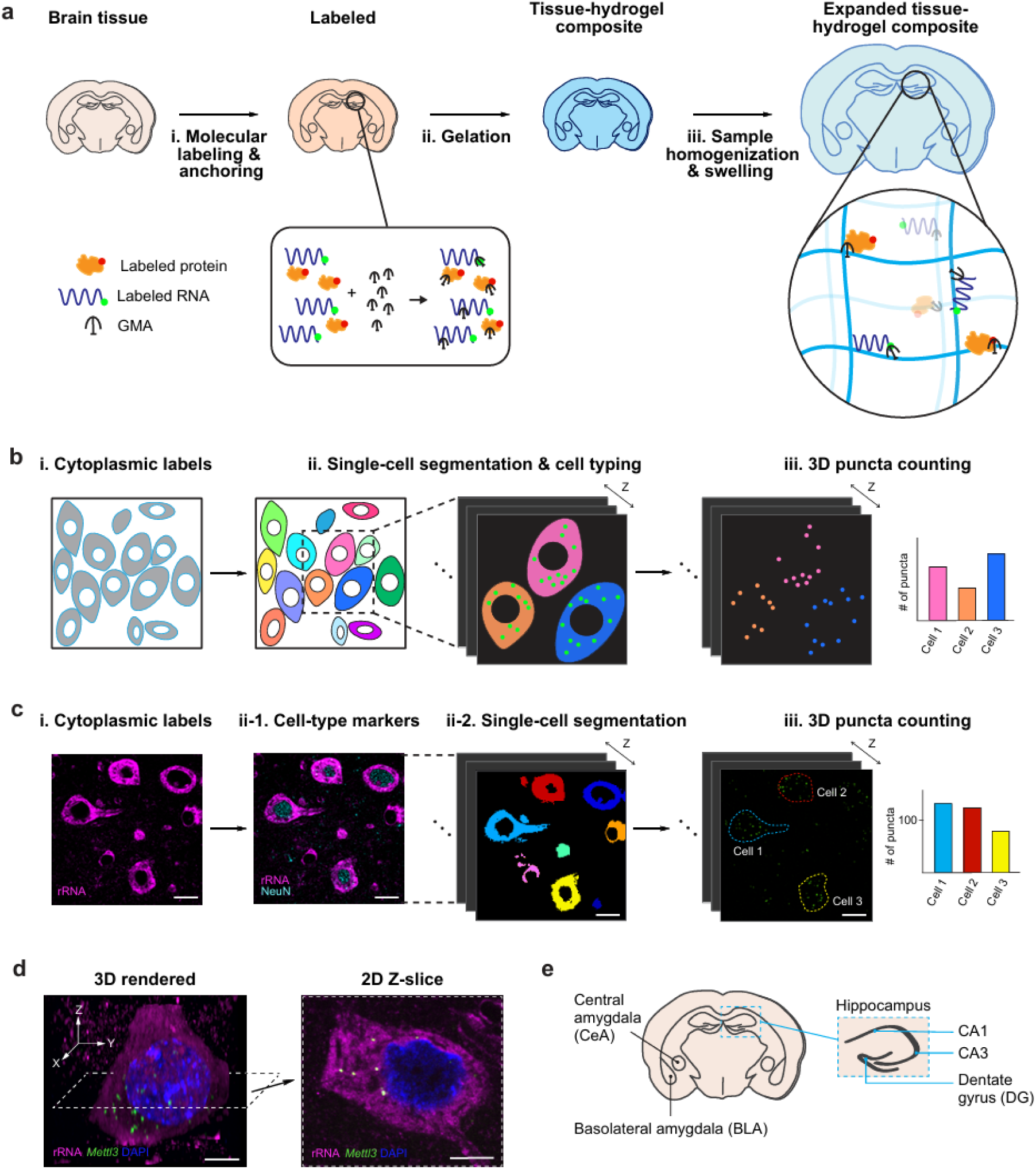
Workflow and validation of RESOLVE-FISH. **(a)** Schematics showing the experimental workflow of RESOLution enhanced Visualization using Expansion-coupled FISH (RESOLVE-FISH). (i) Brain tissue undergoes Hybridization Chain Reaction (HCR)-RNA-FISH and immunostaining to label the target RNAs and proteins, respectively. A multifunctional anchor, glycidyl methacrylate (GMA), is introduced to covalently retain the target biomolecules and labels (rectangular box). (ii) The tissue is embedded in a swellable hydrogel to form a tissue-hydrogel composite. (iii) The tissue is homogenized and expanded, with the target biomolecules and labels retained (circular box). **(b)** Schematics showing the computational and analysis workflow of RESOLVE-FISH. (i) Cytoplasmic labels are used to segment and generate cell masks. (ii) The segmented cells are classified based on the cell-type markers. (iii) Individual RNA puncta are counted in three-dimensions (3D) within each cell mask, enabling spatially-resolved quantification of RNA in single cells across different cell types. **(c)** Fluorescent images from rat basolateral amygdala (BLA) demonstrating the computational and analysis workflow of RESOLVE-FISH. (i) The cytoplasm is labeled using ribosomal RNA (rRNA, magenta). (ii-1) A cell-type marker, NeuN (cyan, neuronal), is used to determine the cell types. (ii-2) Cell segmentation is performed using the cytoplasmic label to generate the cell masks. (iii) RNA puncta (green) are detected and quantified in 3D within each segmented cell, allowing for cell-type-specific quantification of RNA. Scale bars, 10 µm (35 µm). Here and after, scale bars are provided at the pre-expansion scale (with the corresponding post-expansion size indicated in brackets). **(d)** A 3D rendered volumetric image (left) of a neuronal cell from rat BLA and a two-dimensional (2D) Z-slice image (right) of the same cell. The 2D image roughly corresponds to the cross-section indicated by the dashed lines in the 3D rendering. Nuclear label (DAPI, blue), cytoplasmic label (rRNA, magenta), and *Mettl3* RNA puncta (green) are shown. Scale bars, 5 µm (17.5 µm) **(e)** Schematics showing a rat brain coronal section highlighting three brain regions used in this study: the central amygdala (CeA), BLA, and hippocampus. The inset shows a zoomed-in view of the hippocampus (dashed box) highlighting the subregions used in the study: CA1, CA3, and dentate gyrus (DG).

The overall workflow of RESOLVE-FISH starts with fluorescently labeling target molecules, such as mRNAs and proteins of interest, in intact brain tissue (**Fig. 1a**). In this study, we used Hybridization Chain Reaction RNA fluorescence *in situ* hybridization v3.0 (HCR RNA-FISH v3) (Choi et al., 2018), a highly sensitive and quantitative RNA labeling strategy, to label the *Mettl3* mRNA as well as cytoplasmic ribosomal RNA (rRNA). This was followed by an immunostaining step to label cytoplasmic or membrane proteins for cell-typing purposes. We note that this labeling sequence maximizes signal retention and specificity, yielding brighter and more distinct signals compared to alternative orders.

Following the labeling steps, a small-molecule anchor is introduced to link the target molecules and fluorescent labels to a superabsorbent hydrogel matrix. In this study, we used the epoxide-based glycidyl methacrylate (GMA) to covalently anchor both RNAs and proteins to the hydrogel polymer network (Cui et al., 2023). After the anchoring step, the brain tissue is embedded in a superabsorbent hydrogel and enzymatically digested to enable a homogenous expansion. Finally, the gelled brain tissue is expanded multiple-fold to enhance the spatial resolution for the subsequent multi-color fluorescence imaging. This increased resolution allows for spatial delineation of the fluorescently labeled molecules and their downstream quantification at the subcellular level.

For single-cell quantification of target biomolecules, we developed an open-sourced computational pipeline for RESOLVE-FISH, designed to segment individual cells and assign the corresponding fluorescent puncta to the distinct cellular and subcellular compartments (**Fig. 1b, 1c**). First, 3D cell masks are generated using cytoplasmic labels to delineate the mask boundaries, ensuring a robust cell segmentation (**Fig. 1b, i–ii**). For cytoplasmic labeling, a dense fluorescent staining of rRNA or cytoplasmic proteins (e.g., IBA1 for microglia) can be used (**Fig. 1c, i-ii**) (**Methods and Materials**) (Wang et al., 2021). After 3D cell segmentation, the cell-type markers are used to assign the segmented cells to specific cell types (**Fig. 1b, ii**; **Fig. 1c**, **ii**). Finally, 3D quantification of fluorescent puncta, which represent the molecules of interest, is performed for each segmented cells belonging to a specific cell type (**Fig. 1b, iii; Fig. 1c, iii**). For initial validation of its quantitative capability, we applied RESOLVE-FISH to brain (sub)regions known to be related to alcohol addiction—the central amygdala (CeA), basolateral amygdala (BLA), and hippocampus—to assess the transcriptomic changes of *Mettl3* (**Fig. 1d-1e**) (Läck et al., 2007; Geil et al., 2014; Melkumyan and Silberman, 2022). With ∼3.5-fold linear expansion, the final fluorescence volumetric images of the tissue achieved an effective lateral resolution of ∼85 nm, which was sufficient to resolve individual *Mettl3* mRNA puncta within the cytoplasm (**Fig. 1d**, **Supplemental Movie S1**).

### AIE-exposed brain shows cell-type-specific changes in *Mettl3* mRNA across major brain regions in adulthood

To study cell and cell-type-specific AIE-induced changes in the *Mettl3* levels, we applied RESOLVE-FISH to quantify *Mettl3* mRNA across three major cell types—neurons, astrocytes, and microglia—in the CeA of the AIE and Adolescent Intermittent Saline (AIS, control) rats in adulthood. Here, cell-type protein markers (e.g., NeuN for neurons, GFAP for astrocytes, and IBA1 for microglia) were used to delineate each cell population (**Fig. 2a-2c**). Given that our prior study identified no sex-dependent differences in the *Mettl3* mRNA expression following AIE exposure, our experiments were conducted with n = 6 animals per group, with both males and females (n = 3 per sex) (Malovic et al., 2025). As described, the cytoplasmic rRNA and IBA1 signals were used to define the cytoplasmic boundaries of neurons (or astrocytes) and microglia, respectively (**Materials and Methods**), allowing accurate assignments of *Mettl3* mRNA puncta to the cytoplasm of these cells. As a result, we found a significant increase (∼2-fold) in the number of cytoplasmic *Mettl3* mRNA puncta per cell in the CeA neurons of the AIE adult rats [t(10) = -7.35, *p* = 2.78 x 10^-5^, two-sample t-test)] (**Fig. 2d**). In contrast, the *Mettl3* counts in the CeA astrocytes and microglia did not show a significant difference (**Fig. 2e, f**). These findings confirm and extend our previous findings (Malovic et al., 2025), and further suggest that *Mettl3* upregulation in the CeA following AIE exposure is neuron-specific, as no significant changes were observed in microglia or astrocytes.

**Fig. 2:**
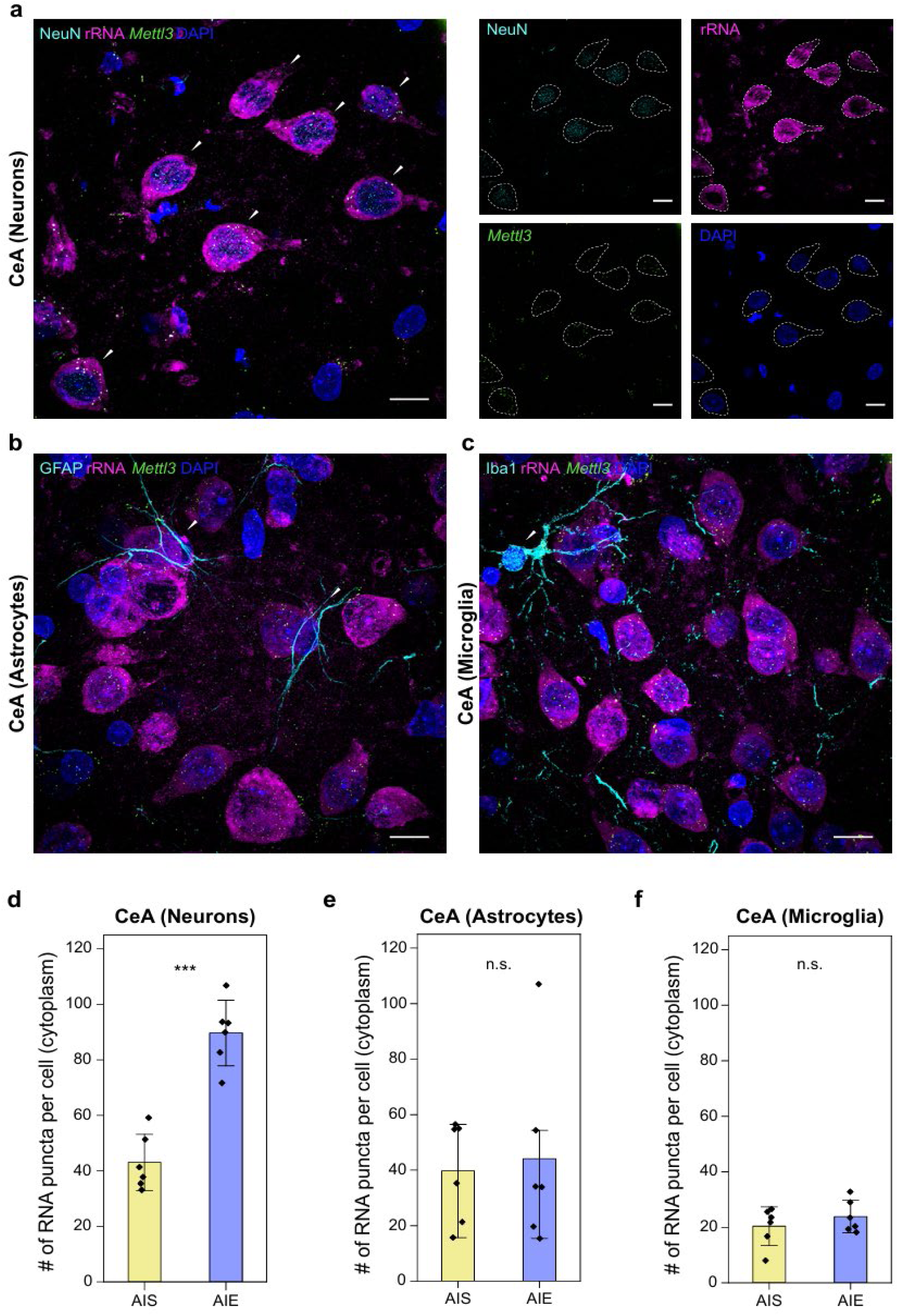
Quantification of *Mettl3* mRNA in the adult central amygdala (CeA) following adolescent alcohol exposure. **(a)** Left: RESOLVE-FISH images showing *Mettl3* mRNA (green) in the CeA of AIE rats. Ribosomal RNA (rRNA, magenta) was used as a cytoplasmic label for single-cell segmentation, and NeuN (cyan) was used to identify neuronal cells. White arrows indicate NeuN-positive cells. Right: Images corresponding to the NeuN, rRNA, *Mettl3* mRNA, and DAPI channels with dashed lines outlining the cell bodies. **(b, c)** RESOLVE-FISH images showing *Mettl3* mRNA (green) in **(b)** GFAP-positive astrocytes (white arrows) and **(c)** IBA1-positive microglia (white arrows) in the CeA of AIE rats. All the images in **(a-c)** are maximum intensity projection (MIP) images (∼6.9 µm in z-range, pre-expansion scale). Scale bars, 10 µm (35 µm). **(d-f)** Cytoplasmic *Mettl3* mRNA puncta per cell across **(d**) neurons (*p* = 2.78 x 10^-5^), **(e)** astrocytes (*p* = 0.790), and **(f)** microglia (*p* = 0.371) in the CeA of AIS and AIE rats (n = 6 per group, male and female, two-sample t-test). In all plots, data are presented as mean ± standard error of the mean (SEM) with individual data points shown as black diamonds.

Next, we examined whether a similar trend can be observed in another amygdalar nucleus. In the BLA, another integral region of the amygdala complex responsible for alcohol exposure-related behaviors (Zhou et al., 2010; Wassum and Izquierdo, 2015), for example, we observed no significant differences in the *Mettl3* levels between the AIE and AIS animals for all three cell types (**Fig. 3**). This brain-region-specific RESOLVE-FISH data suggest that the upregulation of *Mettl3* previously observed via bulk qPCR analysis in the whole amygdala is driven by the CeA (or other amygdalar subregions) rather than the BLA (Malovic et al., 2025). This highlights the high degree of spatial heterogeneity in *Mettl3* regulation across the adult amygdala after AIE exposure.

**Fig. 3:**
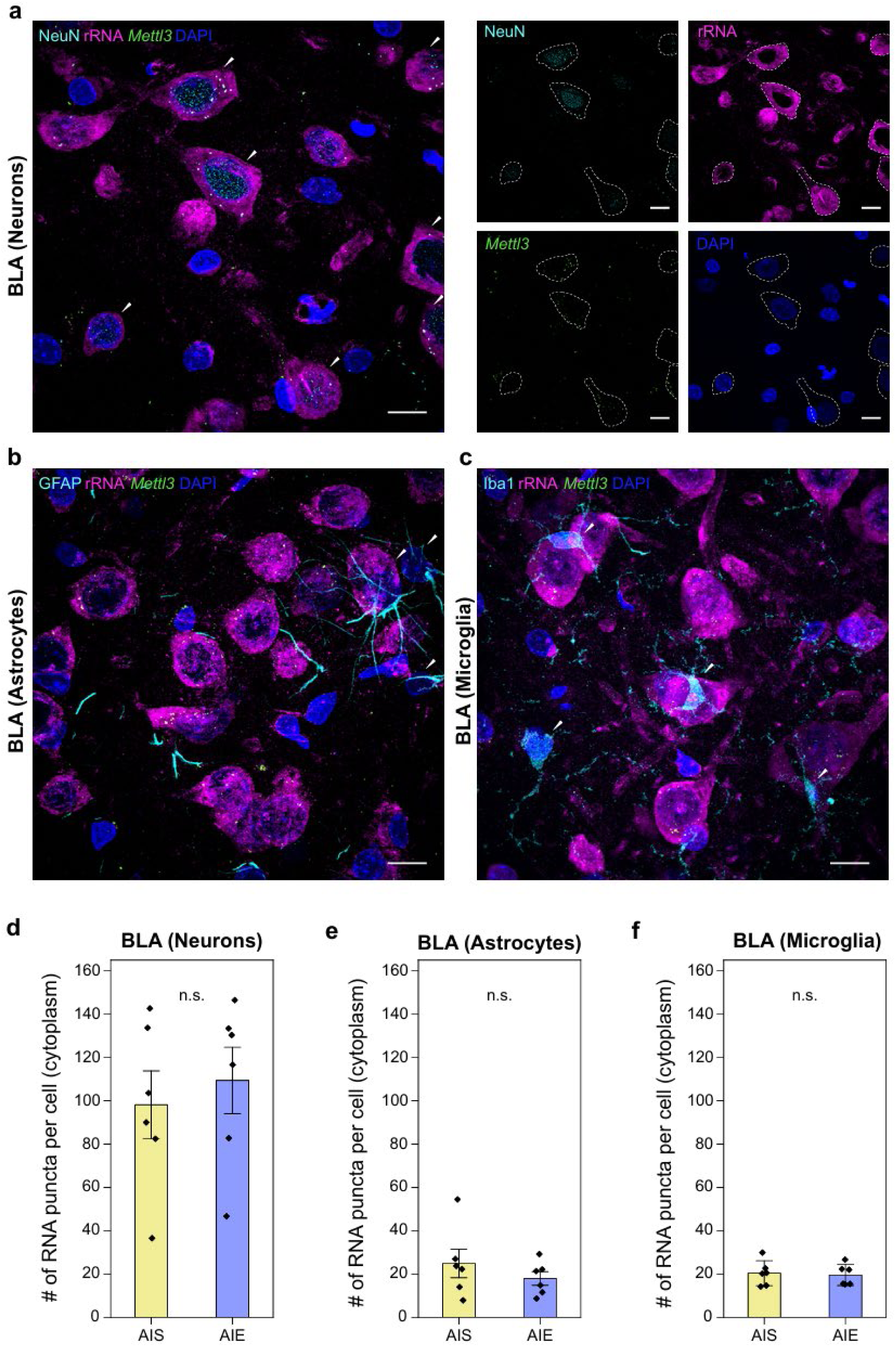
Quantification of *Mettl3* mRNA in the adult basolateral amygdala (BLA) following adolescent alcohol exposure. **(a)** Left: RESOLVE-FISH images showing *Mettl3* mRNA (green) in the BLA of AIE rats. The cytoplasm is labeled by rRNA (magenta), and all neuronal cells (white arrows) are marked by NeuN (cyan). Right: images corresponding to the NeuN, rRNA, *Mettl3* mRNA, and DAPI channels with dashed lines outlining the cell bodies. **(b, c)** RESOLVE-FISH images showing *Mettl3* mRNA (green) in **(b)** GFAP-positive astrocytes (white arrows) and **(c)** IBA1-positive microglia (white arrows) in the BLA of AIE rats. All the images in **(a-c)** are MIP images (∼8.6 µm in z-range, pre-expansion scale). Scale bars, 10 µm (35 µm). **(d-f)** Cytoplasmic *Mettl3* mRNA puncta per cell across **(d)** neurons (*p* = 0.990), **(e)** astrocytes (*p* = 0.371), and **(f)** microglia (*p* = 0.792) in the BLA of AIS and AIE rats (n = 6 per group, male and female, two-sample t-test). In all plots, data are presented as mean ± SEM with individual data points shown as black diamonds.

In contrast to the BLA, we observed a >2-fold increase of *Mettl3* mRNA in neurons within the hippocampus from the AIE animals [t(10) = -3.73, *p* = 0.009, two-sample t-test], exhibiting a finding consistent with that seen in the CeA (**Fig. 4a, 4d**). It is interesting to point out that this effect was not detected in the qPCR study of whole dorsal hippocampal tissue, which reported no changes in the *Mettl3* mRNA level in adulthood (Malovic et al., 2025). Similar to the CeA, this increase in *Mettl3* was not observed in hippocampal astrocytes or microglia (**Fig. 4e, 4f**), again suggesting neuron-specific changes in this brain region. To further investigate the spatial patterns of *Mettl3* expression in the hippocampus, we analyzed the *Mettl3* mRNA counts across hippocampal subregions of CA1, CA3, and dentate gyrus (DG). As a result, both the CA1 and DG neurons exhibited increased *Mettl3* expression in the AIE animals [CA1: t(10) = -3.53, *p* = 0.012, two-sample t-test; DG: t(10) = -3.82, *p* = 0.011, two-sample t-test] (**Supplemental Fig. S1a, S1c**), whereas the CA3 neurons showed no significant differences between the two animal groups (**Supplemental Fig. S1b**). Across all three subregions, *Mettl3* mRNA counts in astrocytes and microglia were not significantly altered by the adolescent alcohol exposure (**Supplemental Fig. S2a-S2c; Supplemental Fig. S3a-S3c**). These results demonstrate that our spatial-resolved and quantitative RESOLVE-FISH pipeline is capable of uncovering neuronal *Mettl3* mRNA increases within the CA1 and DG that are otherwise masked in the bulk qPCR analysis of the whole dorsal hippocampus.

**Fig. 4:**
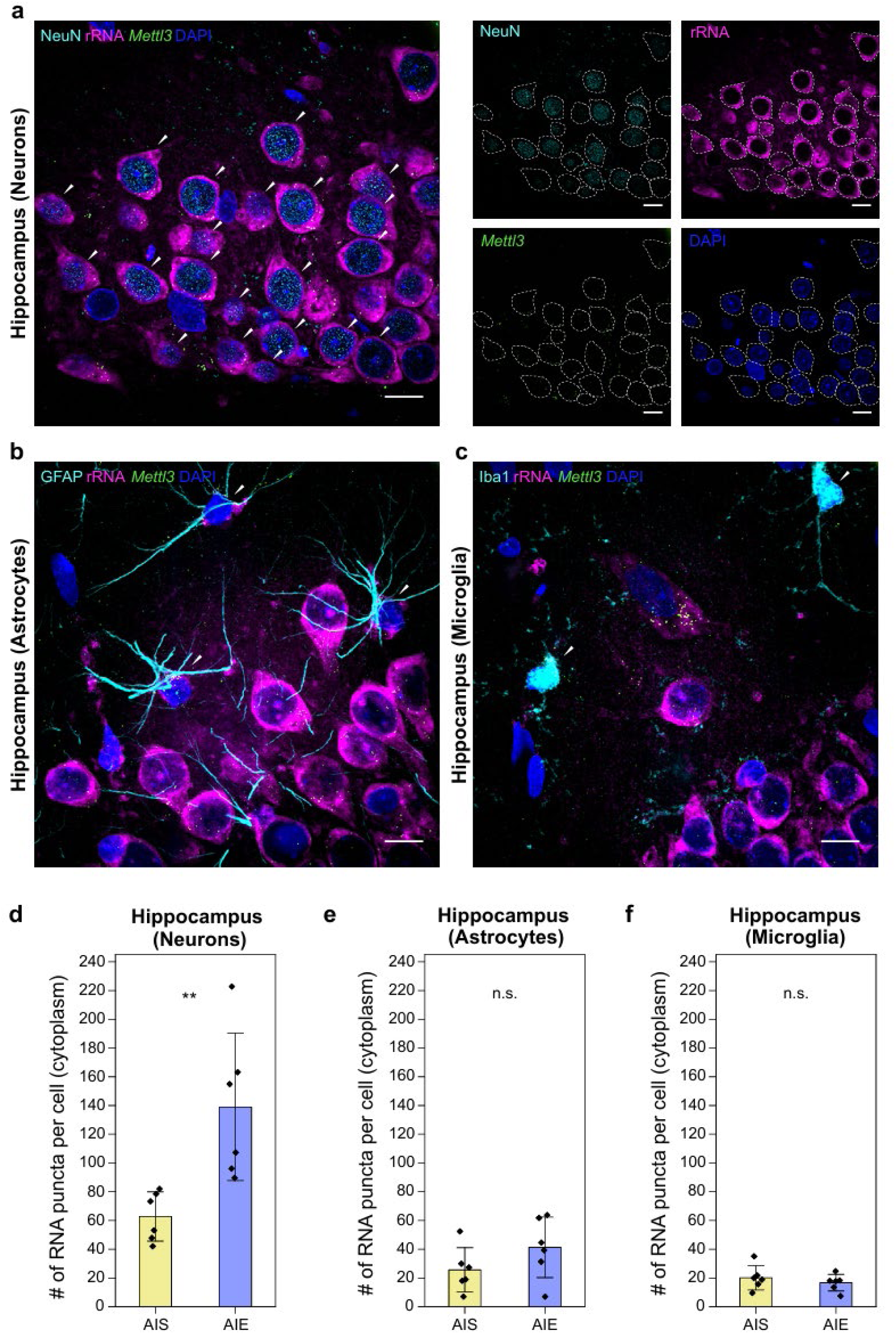
Quantification of *Mettl3* mRNA in the adult hippocampus following adolescent alcohol exposure. **(a)** Left: RESOLVE-FISH images showing *Mettl3* mRNA (green) in the CA1 of AIE rats. The cytoplasm is labeled by rRNA (magenta), and all neuronal cells (white arrows) are marked by NeuN (cyan). Right: images corresponding to the NeuN, rRNA, *Mettl3* mRNA, and DAPI channels, with dashed lines outlining the cell bodies. **(b, c)** RESOLVE-FISH images showing *Mettl3* mRNA (green) in **(b)** GFAP-positive astrocytes (white arrows) in the CA1 and **(c)** IBA1-positive microglia (white arrows) in the DG of AIE rats. All the images in **(a-c)** are MIP images (∼6.9 µm in z-range, pre-expansion scale). Scale bars, 10 µm (35 µm). **(d-f)** Cytoplasmic *Mettl3* mRNA puncta per cell across **(d)** neurons (*p =* 0.009), **(e)** astrocytes (*p* = 0.174), and **(f)** microglia (*p* = 0.421) in the hippocampus of AIS and AIE rats (n = 6 per group, male and female, two-sample t-test). Individual data points are the average of 9 cells, 3 each from the CA1, CA3, and DG. In all plots, data are presented as mean ± SEM with individual data points shown as black diamonds.

### Super-resolved spatial transcriptomics analysis shows cell-type and brain-region-specific differences in *Mettl3* RNA density and subcellular localization

The number of mRNA copies is highly correlated with the cell size, with both mRNA concentration and total copy numbers serving as key indicators of gene expression and resulting protein levels (Edfors et al., 2016; Lin and Amir, 2018). The high resolution and sensitivity of RESOLVE-FISH allowed us to precisely measure the cytoplasmic volume of each cell in correspondence to the number of *Mettl3* puncta measured. As a result, we found that cytoplasmic *Mettl3* mRNA puncta per cell increased across all three brain regions (i.e., in the CeA, BLA, and hippocampus) as the cytoplasmic volume increased (**Fig. 5a-5c**). Grouping by the cell types further revealed that neurons constituted the largest cytoplasmic volumes and exhibited the highest puncta counts, whereas astrocytes and microglia occupied lower ranges for both measures in these brain regions. Importantly, the lines of best fit (shown as the solid lines) were consistently steeper for the AIE animals in the CeA and hippocampus (but not in the BLA), validating our earlier observation of increased *Mettl3* expression in neurons from these regions.

**Fig. 5:**
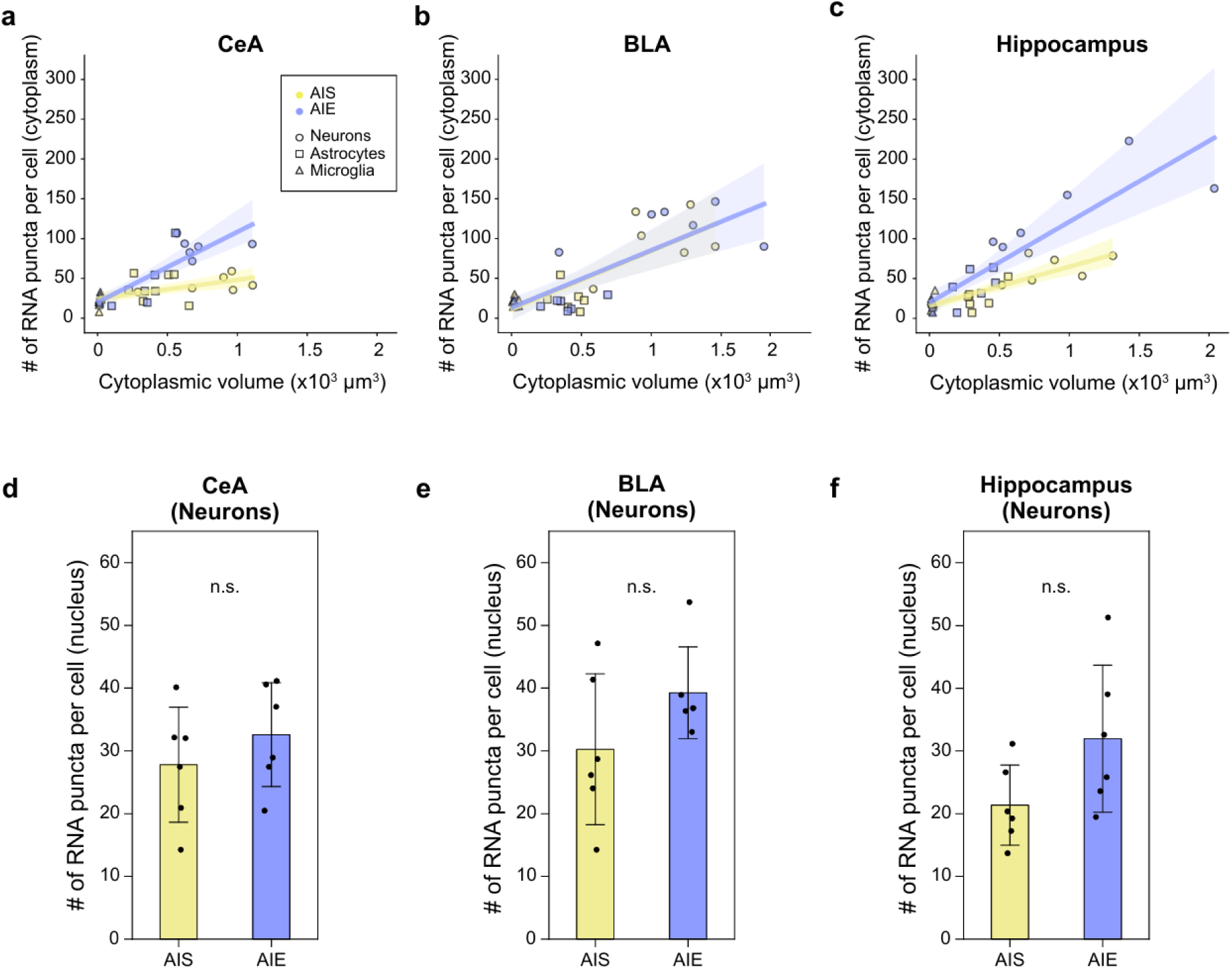
Subcellular quantification and analysis of *Mettl3* RNA across adult brain regions following adolescent alcohol exposure. (a-c) Scatter plots showing the number of cytoplasmic *Mettl3* mRNA puncta and corresponding cytoplasmic volumes in the **(a)** CeA, **(b)** BLA, and **(c)** hippocampus of AIS (yellow) and AIE (violet) rats (n = 6 per group, male and female). Circles, squares, and triangles indicate neurons, astrocytes, and microglia, respectively. Regression lines (solid lines) indicate the trend for the AIS and AIE groups, with the shaded region indicating the 95% confidence interval. **(d-f)** Nucleus-specific quantification of *Mettl3* RNA in neurons in the **(d)** CeA (*p =* 0.366), **(e)** BLA (*p =* 0.155), and **(f)** hippocampus (*p =* 0.089) of AIS and AIE rats (n = 6 per group, male and female, two-sample t-test). Data are presented as mean ± SEM with individual data points shown as black circles, each circle indicating one animal.

To account for inherent differences in the cell size, we further assessed the volumetric density of cytoplasmic *Mettl3* mRNA puncta across the cell types and brain regions studied (**Supplemental Figure S4**). Interestingly, within each animal groups, microglia consistently displayed higher densities than neurons and astrocytes across all regions (**Supplemental Table S3**). This effect was most striking in the CeA, where microglia in the AIE animals exhibited the highest mRNA puncta density across all brain regions and cell types studied (**Supplemental Figure S4b**).

Finally, accurate quantification and comparison of mRNA levels required confining the puncta counting to the cytoplasmic compartment. With its subcellular spatial resolution and high-contrast nuclear labeling, our RESOLVE-FISH pipeline enabled quantification of the *Mettl3* RNA puncta also within the nucleus (**Materials and Methods**). In contrast to the significant differences observed in the cytoplasm, the number of nuclear *Mettl3* RNA puncta did not show a significant difference within neurons for any of the brain regions studied (**Fig. 5d-5f**). A significant difference was then restored for the CeA and hippocampus neurons when we measured the total *Mettl3* RNA puncta in the cell body by combining the nuclear and cytoplasmic RNA puncta (**Supplemental Figure S5**). These results suggest that the observed significant differences are localized specifically to the cytoplasmic mRNA pool rather than other subcellular compartments. For both astrocytes and microglia, the nuclear *Mettl3* RNA counts followed a similar trend to the cytoplasmic results, with no significant differences found between the animal groups (**Supplemental Fig. S6**). Together, these findings indicate that the measured *Mettl3* expression level is greatly dependent on the subcellular compartment from which it is quantified.

## Discussion

We have developed a super-resolution, 3D spatial transcriptomics pipeline, RESOLVE-FISH, to achieve highly sensitive detection of mRNA, concurrent cell-typing, and post-imaging single-cell quantification of the mRNA level. The enhanced spatial resolution as well as the increased intramolecular distances in the expanded tissue allowed for robust quantification of biomolecules of interest, including the target mRNA. When combined with 3D confocal imaging, this “decrowding” effect was essential for the stereological delineation and accurate counting of individual *Mettl3* mRNA puncta within subcellular compartments.

Applying RESOLVE-FISH to brains subjected to AIE exposure, we found that the *Mettl3* mRNA level is selectively increased in a neuron-specific manner in the CeA and certain hippocampal subregions (CA1 and DG), but remains unchanged in astrocytes and microglia as well as in adjacent subregions. This specificity to cell types and brain subregions affirms the link between *Mettl3* mRNA expression and neural circuit plasticity, suggesting that heightened *Mettl3* activity may be functionally relevant to the altered neuronal activity and behavior observed in adulthood after AIE exposure. Supporting this link at the behavioral level, we previously observed that treatment with METTL3 inhibitor at adulthood attenuated anxiety-like behaviors in AIE animals of both sexes (Malovic et al., 2025). Conversely, the lack of *Mettl3* changes in the AIE glial cells suggests that the m6A-mediated regulatory response to AIE exposure is primarily a neuronal phenomenon, likely tied to neuronal plasticity and neurotransmission changes, whereas glial adaptations to AIE exposure may occur through different molecular mechanisms.

Moreover, the observed spatial heterogeneity of *Mettl3* expression in the amygdalar and hippocampal subregions indicates that the cell-type and cell-function-specific epitranscriptomic changes might be important for neuropathogenesis of AIE animals in adulthood. Notably, in the amygdala, the CeA and BLA differ considerably in cellular compositions and functions. The CeA is composed almost entirely of GABAergic neurons, including both interneurons and inhibitory projection neurons, whereas the BLA predominantly contains excitatory glutamatergic principal neurons along with a small fraction of GABAergic interneurons (Jie et al., 2018). Functionally, the CeA serves as the major output nucleus of the amygdala, forwarding inhibitory projections to downstream targets in the brainstem and hypothalamus, whereas the BLA acts as the principal input hub, integrating emotional and associative information (Gilpin et al., 2015; Yang and Wang, 2017).

In addition, the CeA is rich in stress-related neuropeptides, including both anti-stress and pro-stress signaling systems that have been strongly implicated in the alcohol use disorder (AUD) (Koob and Zorrilla, 2010; Sakharkar et al., 2019). In particular, the GABAergic neurons in the CeA are most susceptible to transcriptomic changes following chronic alcohol exposure (Dilly et al., 2022). Our findings extend this framework by showing enduring epitranscriptomic changes in the CeA (Funk et al., 2006; Sommer et al., 2008). Furthermore, the difference in the *Mettl3* level changes between the CeA and BLA may reflect region-specific demands for RNA methylation-dependent gene regulation, with CeA inhibitory neurons and BLA excitatory neurons having distinct needs for RNA regulation to mediate their roles in emotional processing such as anxiety-like behaviors after early-life or adult alcohol exposure.

In the hippocampus, the CA1 and CA3 are composed primarily of excitatory pyramidal neurons, whereas the DG contains densely packed granule cells and is one of the few brain regions where adult neurogenesis occurs (Mandyam, 2013).

Functionally, the CA1 acts as the main output of the hippocampus, relaying processed information to cortical areas (Valenzuela and Morton, 2014). The CA3 functions as an autoassociative network important for memory encoding and retrieval, and the DG contributes to pattern separation by transforming similar inputs into distinct representations before transmitting them to the CA3 (Bakker et al., 2008).

As previously reported, both the DG and CA1 subregions are highly plastic and potentially more sensitive to alcohol (Chen et al., 2013; Ramachandran et al., 2015; Vetreno et al., 2018). For example, AIE exposure is known to suppress adult neurogenesis in the DG (Vetreno and Crews, 2015; Sakharkar et al., 2016), induce granule cell loss, as well as impair synaptic plasticity and cause neuronal loss in the CA1 (Risher et al., 2015), leading to disrupted hippocampal function (Nixon and Crews, 2002; Nalberczak-Skóra et al., 2023). Again, the selective upregulation of *Mettl3* mRNA in the CA1 and DG neurons may represent a compensatory response aimed at maintaining synaptic integrity and neurogenic capacity under conditions of alcohol-induced stress, as METTL3 is required for normal neurogenesis and cognitive function in adult (Yoon et al., 2017). Interestingly, the CA3 appears more resilient given this compensatory hypothesis, while the CA1 and DG are more sensitive and actively trying to adapt through m6A RNA modification. These findings suggest that not all hippocampal circuits are equally vulnerable to AIE, and that the modulation of key epitranscriptomic switches such as *Mettl3* is highly heterogeneous.

Extending beyond subregional differences, our study highlights that cell type and subcellular compartmentalization are critical determinants of *Mettl3* regulation. From our study, a significantly higher *Mettl3* mRNA puncta density was observed in microglia regardless of the brain region or alcohol exposure condition. This observation may be attributed to phagocytosed materials as it has been previously revealed by RNA-seq that a major portion of the transcripts identified in microglia are expressed by other cell types in the brain, including the neurons (Solga et al., 2015). Subcellular distribution is another factor contributing to the cell-type-specific, spatially heterogeneous expression of *Mettl3*. Because the observed significant increase in the *Mettl3* mRNA in AIE neurons was specific to cytoplasm (in the CeA and hippocampus) and absent in the nucleus, it is essential to confine the mRNA quantification to particular subcellular compartments for biologically meaningful comparisons.

We recently reported that *Mettl3* mRNA levels are increased in the adult CeA in the putative neuronal populations after AIE exposure (Malovic et al., 2025). This finding was further associated with the observed increase in the protein levels of METTL3 and m6A levels in the CeA (Malovic et al., 2025). Our RESOLVE-FISH data thus extend these findings by revealing that *Mettl3* upregulation is cytoplasmic and restricted to neurons within the CeA, as well as the hippocampal CA1 and DG, while remaining absent in astrocytes and microglia across these regions. Consequently, this study addresses a critical knowledge gap in the field by validating that *Mettl3* upregulation is indeed a neuronal effect.

Continuing efforts are needed to adapt the RESOLVE-FISH pipeline for precise quantification of broader epitranscriptomic machinery, including m6A writers, erasers, readers, as well as their downstream mRNA and protein targets. For high-abundance mRNA species, a larger tissue expansion factor may be necessary to resolve individual mRNA puncta. This could be achieved by utilizing superabsorbent hydrogel chemistries capable of ∼10-20-fold linear expansion (Truckenbrodt et al., 2018; Damstra et al., 2022; Klimas et al., 2023; Wang et al., 2024). For detection and quantification of multiple mRNA targets, a multi-round hybridization or a barcoding strategy could be adopted (Chen et al., 2016; Wang et al., 2021). Additionally, the integration of oligo-based barcoding and signal amplification would enable the concurrent detection and quantification of mRNA and the corresponding proteins (Black et al., 2021; Gandin et al., 2025). Ultimately, coupling RESOLVE-FISH with combinatorial labeling or in situ sequencing could offer a pathway toward transcriptome-wide mRNA quantification (Wang et al., 2018; Alon et al., 2021).

## Author Contributions

R.G. and S.C.P. designed research; J.T., E.M., A.I., and H.Z. performed research; J.T., A.I., and B.A. analyzed data; J.T., R.G., and S.C.P. wrote the first draft of the paper; All authors edited the paper.

## Acknowledgement

This work was supported by National Institute on Alcohol Abuse and Alcoholism grants (UO1AA019971, U24AA024605 [Neurobiology of Adolescent Drinking in Adulthood (NADIA) project], and P50AA022538) to S.C.P., the Department of Veterans Affairs (Senior Research Career Scientist award, IK6BX006030, to S.C.P.), and National Institutes of Health grants (pilot program within P50AA022538, DP2MH136390) to R.G. E.M. was supported by the post-doctoral fellowship from T32 AA026577.

