## Supplemental Information for "Super-resolved, three-dimensional spatial transcriptomics reveals cell-type and brain-region-specific modulation of key epitranscriptomic switches following adolescent alcohol exposure"

**Supplemental Tables**

**Table S1: Primary antibodies used in this study.**

| Host | Target | Vendor | Cat. No. |
|---|---|---|---|
| Guinea pig | NeuN | Synaptic Systems | 266 004 |
| Guinea pig | GFAP | Synaptic Systems | 173 308 |
| Chicken | Iba1 | Synaptic Systems | 234 009 |

**Table S2: Secondary antibodies used in this study.**

| Host | Target | Conjugation | Vendor | Cat. No. | Custom conjugation to |
|---|---|---|---|---|---|
| Goat | Guinea pig | Unconjugated | ThermoFisher | A18771 | 647 NHS dye |
| Goat | Chicken | Unconjugated | ThermoFisher | A16056 | 647 NHS dye |

**Table S3: Two-way ANOVA test performed for pair-wise comparison of cytoplasmic puncta density across cell-types and brain regions.**

| | Group 1 | Group 2 | Mean diff. | p-value |
|---|---|---|---|---|
| **AIS** | Astrocytes_BLA | Microglia_BLA | 1.0985 | < 0.0001 |
| | Astrocytes_BLA | Microglia_CeA | 1.0473 | < 0.0001 |
| | Astrocytes_BLA | Microglia_Hippocampus | 0.928 | 0.0005 |
| | Astrocytes_CeA | Microglia_BLA | 1.0527 | < 0.0001 |
| | Astrocytes_CeA | Microglia_CeA | 1.0015 | 0.0001 |
| | Astrocytes_CeA | Microglia_Hippocampus | 0.8822 | 0.013 |
| | Astrocytes_Hippocampus | Microglia_BLA | 1.0938 | < 0.0001 |
| | Astrocytes_Hippocampus | Microglia_CeA | 1.0426 | < 0.0001 |
| | Astrocytes_Hippocampus | Microglia_Hippocampus | 0.9233 | 0.0006 |
| | Microglia_BLA | Neurons_BLA | -1.0707 | < 0.0001 |
| | Microglia_BLA | Neurons_CeA | -1.1043 | < 0.0001 |
| | Microglia_BLA | Neurons_Hippocampus | -1.0894 | < 0.0001 |
| | Microglia_CeA | Neurons_BLA | -1.0195 | 0.0001 |
| | Microglia_CeA | Neurons_CeA | -1.0531 | < 0.0001 |
| | Microglia_CeA | Neurons_Hippocampus | -1.0382 | < 0.0001 |

| | | | | |
|---|---|---|---|---|
| | Microglia_Hippocampus | Neurons_BLA | -0.9002 | 0.0009 |
| | Microglia_Hippocampus | Neurons_CeA | -0.9338 | 0.0005 |
| | Microglia_Hippocampus | Neurons_Hippocampus | -0.9189 | 0.0006 |

| | | | | |
|---|---|---|---|---|
| **AIE** | Astrocytes_BLA | Microglia_BLA | 1.0098 | 0.0001 |
| | Astrocytes_BLA | Microglia_CeA | 1.6288 | < 0.0001 |
| | Astrocytes_BLA | Microglia_Hippocampus | 0.8576 | 0.0021 |
| | Astrocytes_CeA | Microglia_BLA | 0.9392 | 0.0004 |
| | Astrocytes_CeA | Microglia_CeA | 1.5582 | < 0.0001 |
| | Astrocytes_CeA | Microglia_Hippocampus | 0.787 | 0.079 |
| | Astrocytes_Hippocampus | Microglia_BLA | 0.9247 | 0.0005 |
| | Astrocytes_Hippocampus | Microglia_CeA | 1.5436 | < 0.0001 |
| | Astrocytes_Hippocampus | Microglia_Hippocampus | 0.7724 | 0.0103 |
| | Microglia_BLA | Neurons_BLA | -0.9356 | 0.0004 |
| | Microglia_BLA | Neurons_CeA | -0.9285 | 0.0005 |
| | Microglia_BLA | Neurons_Hippocampus | -0.9017 | 0.0009 |
| | Microglia_CeA | Microglia_Hippocampus | -0.7712 | 0.0105 |
| | Microglia_CeA | Neurons_BLA | -1.5546 | < 0.0001 |
| | Microglia_CeA | Neurons_CeA | -1.5475 | < 0.0001 |
| | Microglia_CeA | Neurons_Hippocampus | -1.5206 | < 0.0001 |
| | Microglia_Hippocampus | Neurons_BLA | -0.7834 | 0.0085 |
| | Microglia_Hippocampus | Neurons_CeA | -0.7763 | 0.0096 |
| | Microglia_Hippocampus | Neurons_Hippocampus | -0.7494 | 0.0154 |

#### Supplemental Figures

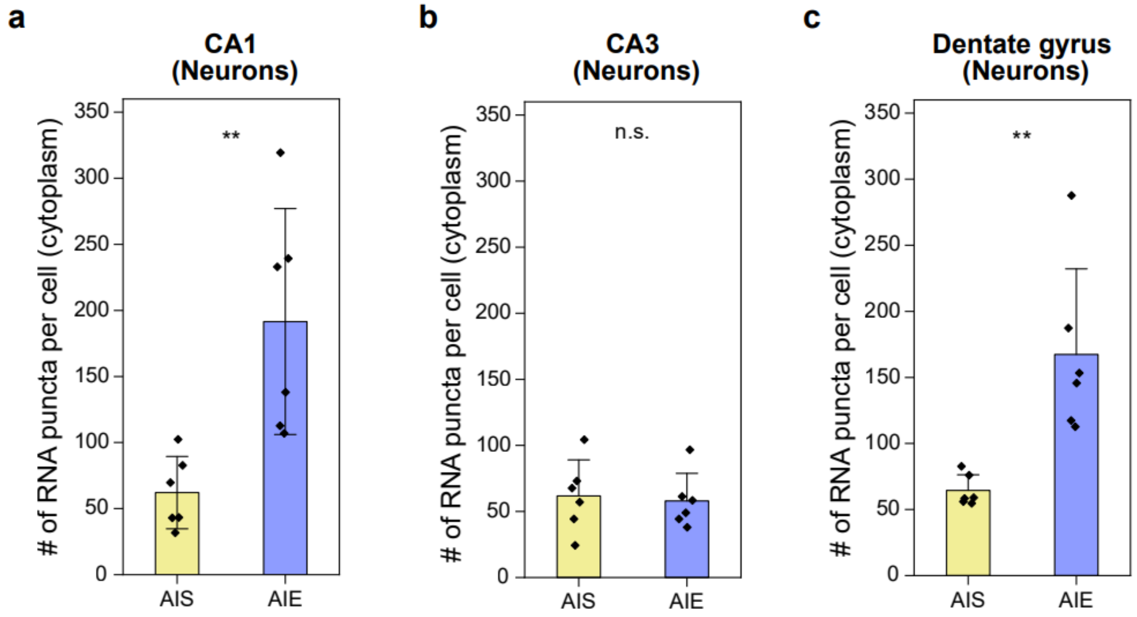


**Fig. S1: RESOLVE-FISH quantification of cytoplasmic *Mettl3* mRNA puncta in neurons of distinct adult hippocampal subregions following adolescent alcohol exposure.** Cytoplasmic *Mettl3* mRNA puncta per cell across neurons in hippocampal subregions of **(a)** CA1 ($p$ = 0.012), **(b)** CA3 ($p$ = 0.790), and **(c)** dentate gyrus ($p$ = 0.011) of AIS and AIE rats (n = 6 per group, male and female, two-sample t-test). Individual data points are the average of 3 cells each from the CA1, CA3, and DG, respectively. In all plots, data are presented as mean ± standard error of the mean (SEM) with individual data points shown as black diamonds.

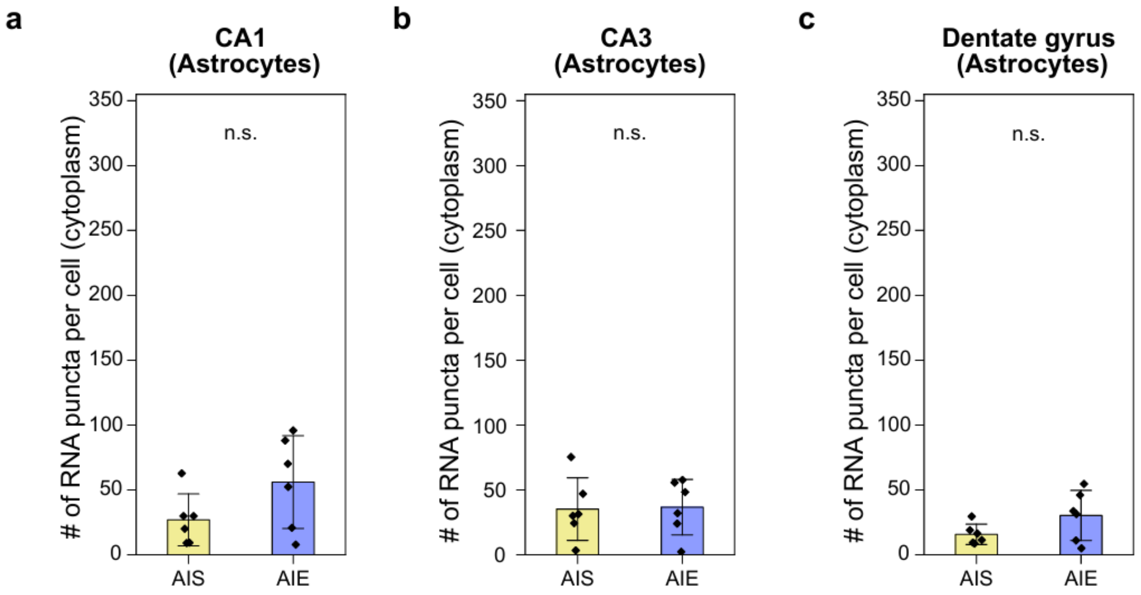


**Fig. S2: RESOLVE-FISH quantification of cytoplasmic *Mettl3* mRNA puncta in astrocytes of distinct adult hippocampal subregions following adolescent alcohol exposure.** Cytoplasmic *Mettl3* mRNA puncta per cell across astrocytes in hippocampal subregions of **(a)** CA1 ($p$ = 0.121), **(b)** CA3 ($p$ = 0.915), and **(c)** dentate gyrus ($p$ = 0.133) of AIS and AIE rats (n = 6 per group, male and female, two-sample t-test). Individual data points are the average of 3 cells each from the CA1, CA3, and DG, respectively. In all plots, data are presented as mean ± SEM with individual data points shown as black diamonds.

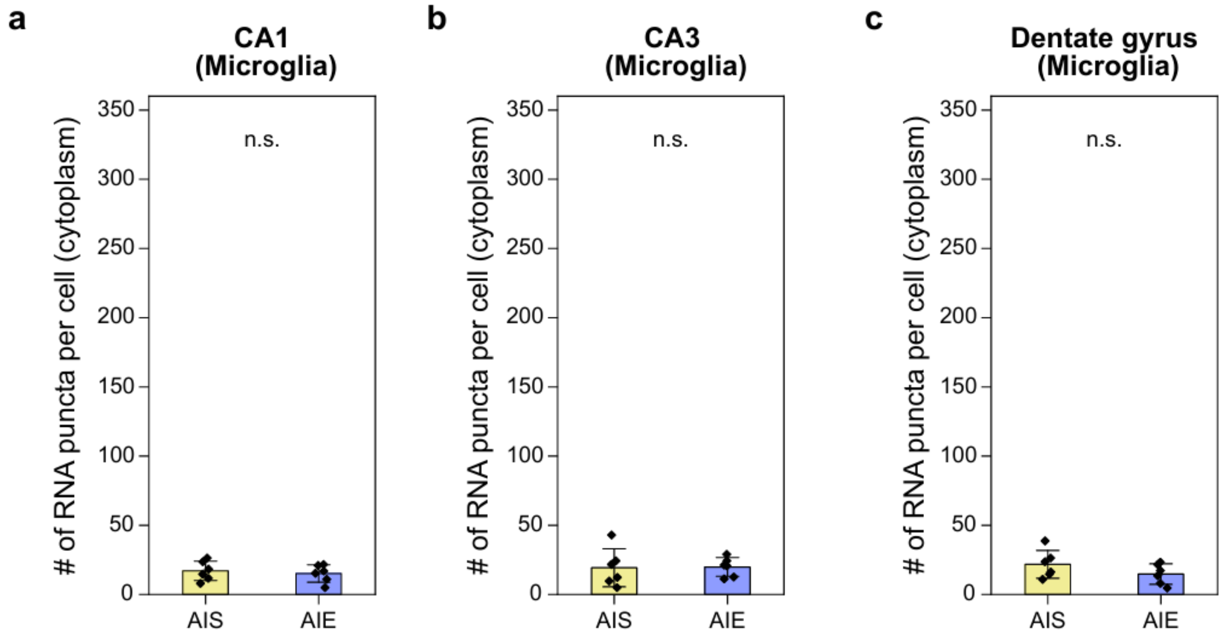


**Fig. S3: RESOLVE-FISH quantification of cytoplasmic *Mettl3 m*RNA puncta in microglia of distinct adult hippocampal subregions following adolescent alcohol exposure.** Cytoplasmic *Mettl3 m*RNA puncta per cell across microglia in hippocampal subregions of **(a)** CA1 ($p$ = 0.626), **(b)** CA3 ($p$ = 0.931), and **(c)** dentate gyrus ($p$ = 0.203) of AIS and AIE rats (n = 6 per group, male and female, two-sample t-test). Individual data points are the average of 3 cells each from the CA1, CA3, and DG, respectively. In all plots, data are presented as mean ± SEM with individual data points shown as black diamonds.

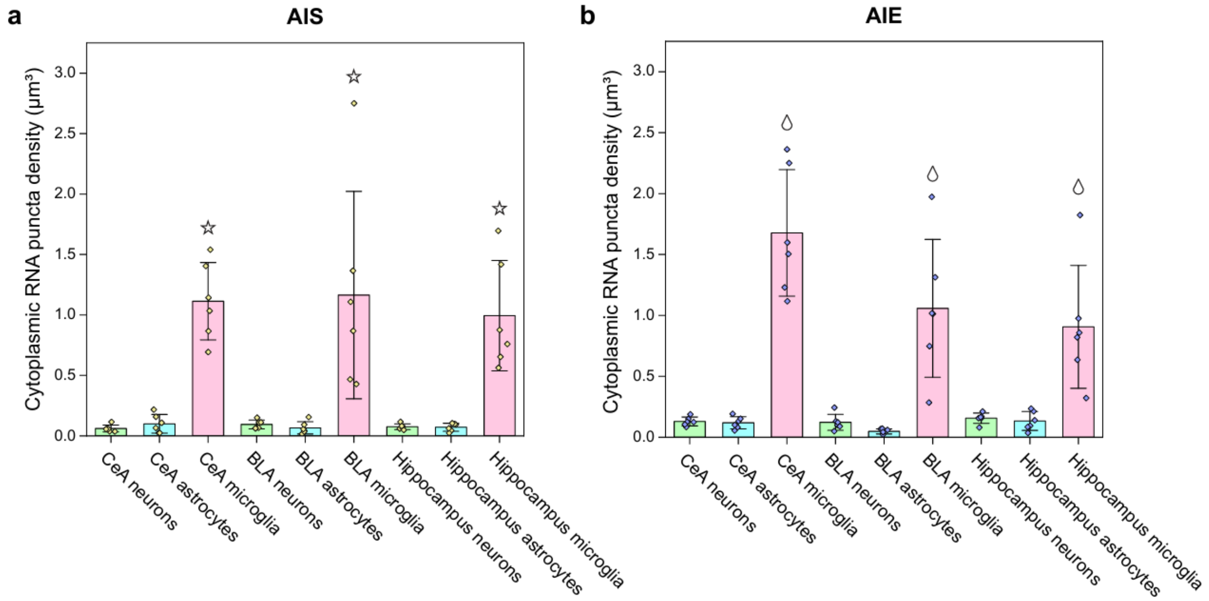


**Fig. S4: RESOLVE-FISH quantification of volumetric density of cytoplasmic *Mettl3* RNA puncta across adult brain regions and cell types following adolescent alcohol exposure.** Volumetric density of cytoplasmic *Mettl3* mRNA were measured in neurons (green), astrocytes (blue), and microglia (pink) in the CeA, BLA, and hippocampus of (a) AIS (yellow diamonds) and (b) AIE (violet diamonds) rats. Stars (in the AIS plot) and droplets (in the AIE plot) indicate significant differences between the microglia density and that for other cell types across all brain regions. Three-way ANOVA was performed for statistical testing with a post hoc Tukey test for pairwise comparisons (with p-values and associated statistics shown in **Table S3**).

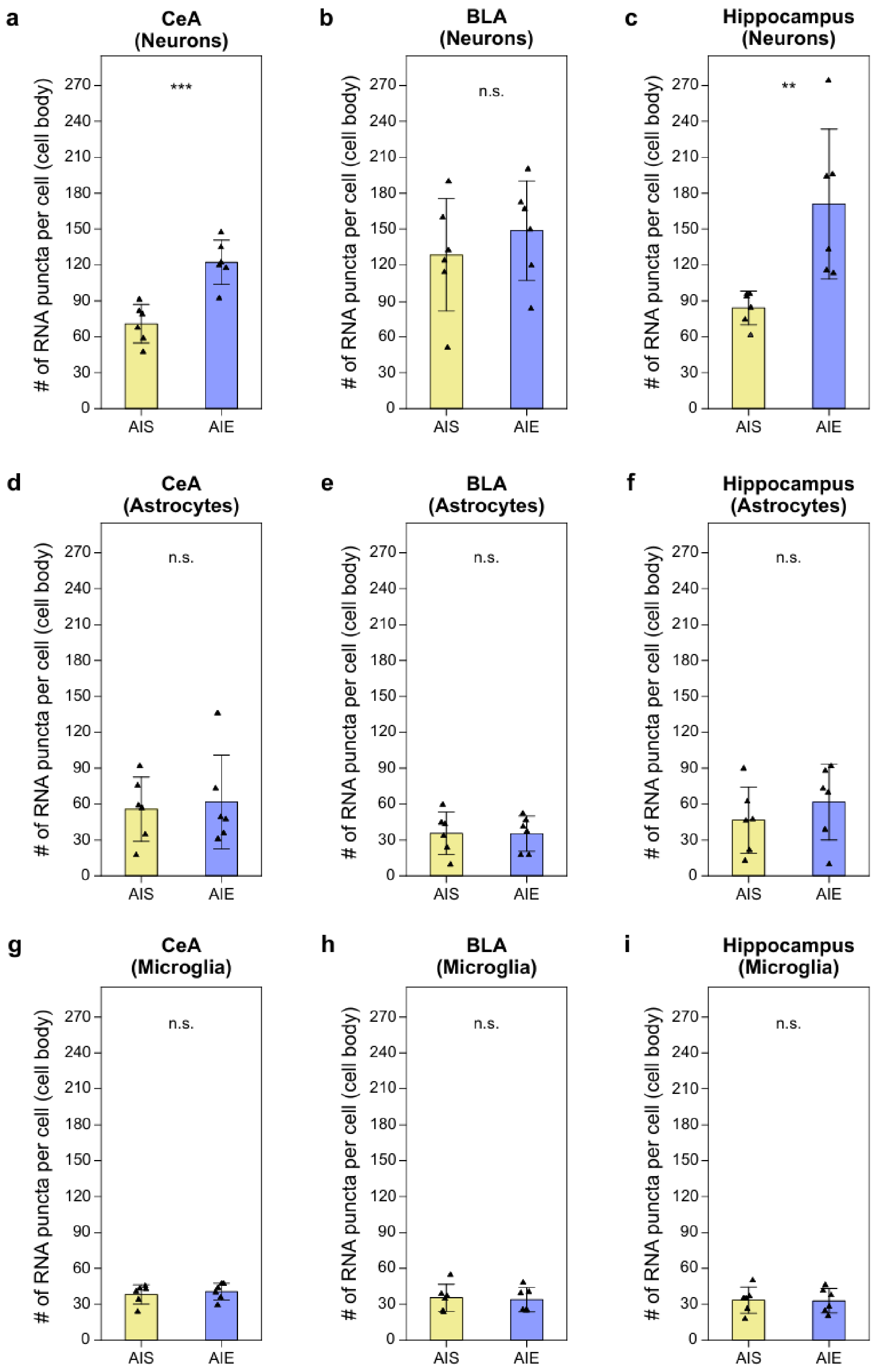

a
CeA (Neurons)
***
b
BLA (Neurons)
n.s.
c
Hippocampus (Neurons)
**
d
CeA (Astrocytes)
n.s.
e
BLA (Astrocytes)
n.s.
f
Hippocampus (Astrocytes)
n.s.
g
CeA (Microglia)
n.s.
h
BLA (Microglia)
n.s.
i
Hippocampus (Microglia)
n.s.
### of RNA puncta per cell (cell body)
0
30
60
90
120
150
180
210
240
270
AIS
AIE

**Fig. S5: RESOLVE-FISH quantification of *Mettl3* RNA puncta in the cell body of different cell types across the adult brain regions following adolescent alcohol exposure.** Total *Mettl3* RNA puncta in the cell body (the sum of nuclear and cytoplasmic puncta) per cell across neurons in the **(a)** CeA ($p$ = 4.791x$10^{-4}$), **(b)** BLA ($p$ = 0.448), and **(c)** hippocampus ($p$ = 0.018), astrocytes in the **(d)** CeA ($p$ = 0.757), **(e)** BLA ($p$ = 0.968), and **(f)** hippocampus ($p$ = 0.399), and microglia in the **(g)** CeA ($p$ = 0.615), **(h)** BLA ($p$ = 0.804), and **(i)** hippocampus ($p$ = 0.934) of AIS and AIE rats (n = 6 per group, male and female, two-sample t-test). In all plots, data are presented as mean ± SEM with individual data points shown as black triangles.

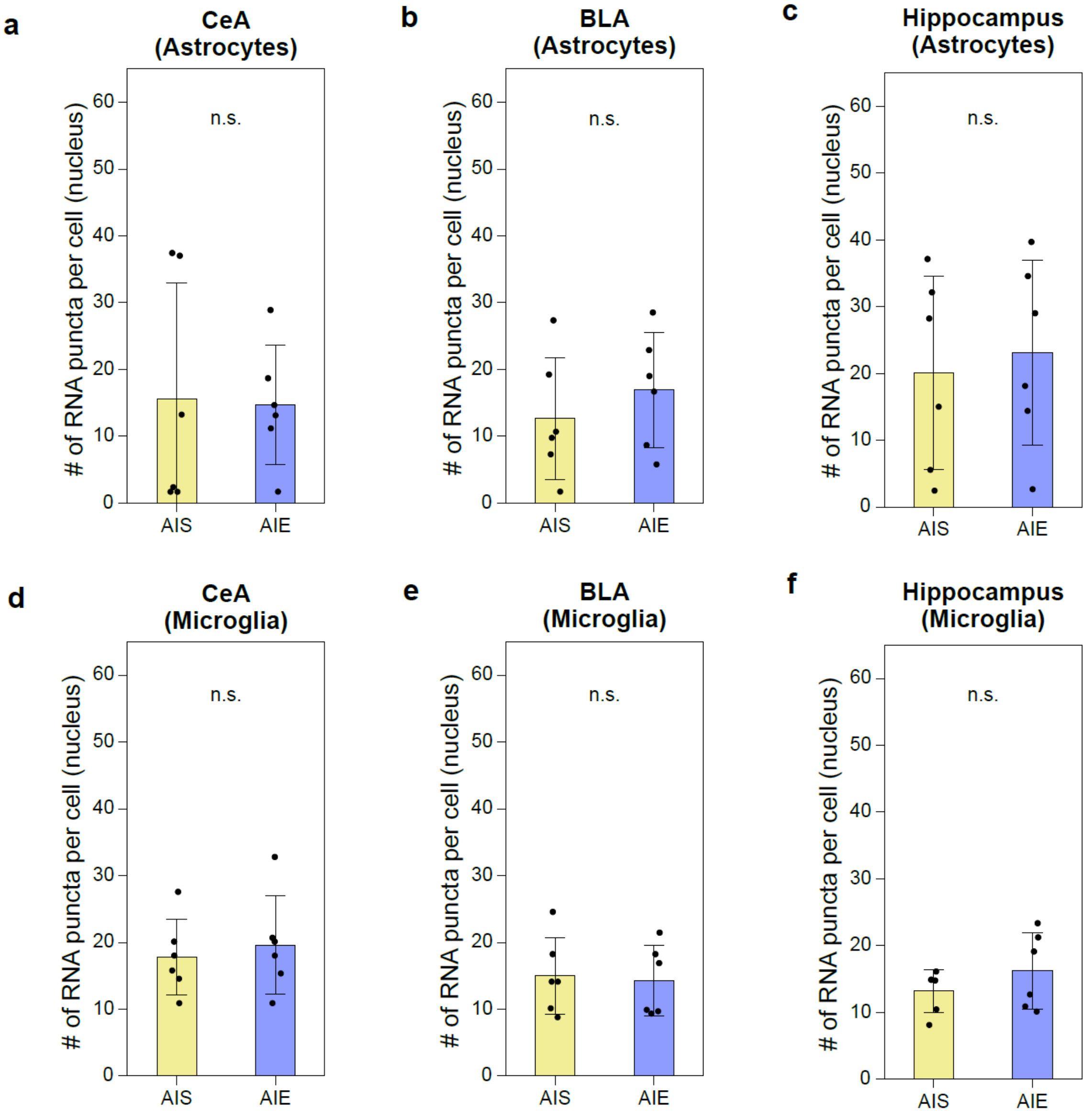


**Fig. S6: RESOLVE-FISH quantification of nuclear *Mettl3* RNA puncta of glial cells across the adult brain regions following adolescent alcohol exposure.** Nuclear *Mettl3* RNA puncta per cell across astrocytes in the **(a)** CeA ($p$ = 0.916), **(b)** BLA ($p$ = 0.425), and **(c)** hippocampus ($p$ = 0.723), and across microglia in the **(d)** CeA ($p$ = 0.644), **(e)** BLA ($p$ = 0.821), and **(f)** hippocampus ($p$ = 0.290) of AIS and AIE rats (n = 6 per group, male and female) (two-sample t-test). In all plots, data are presented as mean ± SEM with individual data points shown as black circles.

**Supplemental Movies**

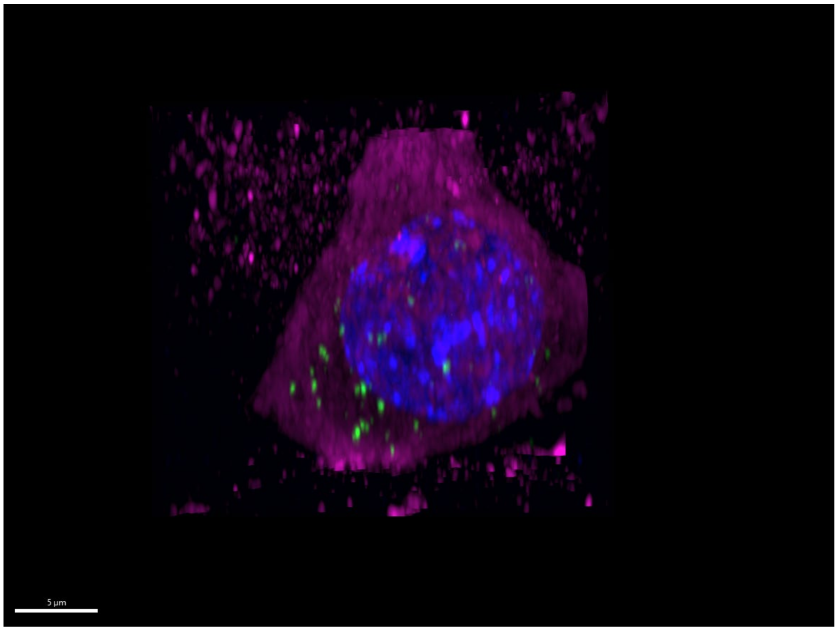


**Movie S1: Three-dimensional (3D) visualization of a neuronal cell imaged with RESOLVE-FISH.** The movie displays a 3D rendering of a neuronal cell reconstructed from an image Z-stack (~21 µm in Z-range, pre-expansion scale), illustrating the spatial distribution of individual RNA puncta within the cytoplasm relative to the nuclear compartment. Cytoplasmic label (rRNA) is shown in magenta, the nucleus (DAPI) in blue, and *Mettl3* mRNA puncta in green. Scale bar, 5 µm (17.5 µm). The scale bar is provided at the pre-expansion scale (with the corresponding post-expansion size indicated in brackets).
